# Tomato spotted wilt virus benefits its thrips vector by modulating metabolic and plant defense pathways in tomato

**DOI:** 10.1101/2020.08.10.244665

**Authors:** Punya Nachappa, Jean Challacombe, David C. Margolies, James R. Nechols, Anna E. Whitfield, Dorith Rotenberg

## Abstract

Several plant viruses modulate vector fitness and behavior in ways that may enhance virus transmission. Previous studies have documented indirect, plant-mediated effects of tomato spotted wilt virus (TSWV) infection on the fecundity, growth and survival of its principal thrips vector, *Frankliniella occidentalis*, the western flower thrips. In this study, we conducted thrips performance and preference experiments combined with plant gene expression, phytohormone and total free amino acid analyses to identify tomato host responses to single and dual challenge with TSWV and *F. occidentalis*, compared to *F. occidentalis* alone, to address the question: do systemically-infected, symptomatic tomato plants modulate primary metabolic (photosynthesis and related physiological functions) and defense-related pathways to culminate into a more favorable environment for the vector. In a greenhouse setting, we documented a significant increase in the number of offspring produced by *F. occidentalis* on TSWV-infected tomato plants compared to mock-inoculated plants, and in choice test assays, females exhibited enhanced settling on TSWV-infected leaves. Microarray analysis combined with phytohormone signaling pathway analysis revealed that TSWV infection, regardless of thrips activity, robustly upregulated salicylic acid (SA) synthesis and downstream defense signaling pathway genes typically known to be associated with execution of defense against pathogens. TSWV alone downregulated a few jasmonic acid (JA)-responsive, anti-herbivore defense genes, however these were limited to wound-induced proteinase inhibitors. While this may indicate a subtle SA-JA antagonistic cross-talk in response to the virus, abscisic acid (ABA, upregulated) and auxin pathways (downregulated) were also perturbed by TSWV infection, regardless of *F. occidentalis* colonization, and may play roles in coordinating and dampening defense against the vector on infected plants. *Frankliniella occidentalis* alone triggered JA and ET pathways, phytohormones that have been reported to work cooperatively to enhance induced resistance to microbes and herbivores; however, on infected plants, ET remained unperturbed by the thrips vector. TSWV infection, alone or in combination with thrips, suppressed genes associated with photosynthesis and chloroplast function thereby significantly impacting primary metabolism of the host plant, and hierarchical cluster analysis and network analysis revealed that many of these genes were co-regulated with phytohormone defense signaling genes. Virus infection also altered genes related to cell wall organization which may render plants more susceptible to the penetration of thrips mouthparts. Lastly, TSWV infection increased expression of genes related to protein synthesis and degradation which is reflected in the increased total free amino acid content in virus-infected plants that harbored higher thrips populations. These results suggest coordinated gene networks that regulate plant primary metabolism and defense responses rendering virus-infected plants more conducive host for vectors, a relationship that is beneficial to the vector and the virus when considered within the context of the complex transmission biology of TSWV. To our knowledge this is the first study to identify global transcriptional networks that underlie the TSWV-thrips interaction as compared to a single mechanistic approach. Findings of this study increase our fundamental knowledge of host plant-virus-vector interactions and identifies underlying mechanisms of induced host susceptibility to the insect vector.

## INTRODUCTION

Most plant-pathogenic viruses depend exclusively on insects for transmission to plants (Hogenhout et al., 2008). It would therefore seem to be evolutionarily advantageous for a plant virus to modify vector behavior and performance in ways that enhance its likelihood of acquisition and dissemination in the landscape (Reviewed in (Eigenbrode et al., 2018). Indeed, there are cases that demonstrate the direct effect of some plant viruses to alter vector performance. For example, acquisition of barley yellow dwarf virus (BYDV) altered aphid vector preference for non-infected compared to BYDV-infected plants (Ingwell et al., 2012) and acquisition of tomato mottle virus increased oviposition of its whitefly vector (McKenzie, 2002). Indirect or plant-mediated effects due to virus infection also play a role in vector-plant interactions. For example, plants infected with potato leafroll virus and BYDV were not only more attractive to aphid vectors, but also increased nutrient quality that enhanced vector survival and fecundity (Eigenbrode et al., 2002; Jimenez-Martinez et al., 2004; Ngumbi et al., 2007). These studies highlight the widespread occurrence of vector manipulation by plant viruses across several pathosystems for aphids (Fereres et al., 1989; Blua et al., 1994; Eigenbrode et al., 2002; Mauck et al., 2010), whiteflies (Guo et al., 2010), and thrips (Carter, 1939; Maris et al., 2004; Belliure et al., 2005; Abe et al., 2012).

Among thrips-transmitted tospoviruses, *Tomato spotted wilt orthotospovirus* (order *Bunyavirales*, family *Tospoviridae*, genus *Orthotospovirus*) is ranked among the top 10 most economically important plant viruses worldwide (Rybicki, 2015). The virus is transmitted in a circulative-persistent manner exclusively by thrips, the most efficient of which is the western flower thrips, *Frankliniella occidentalis* (Pergande) (Whitfield et al., 2005; Whitfield et al., 2015; Rotenberg and Whitfield, 2018). The thrips-tospovirus relationship is unique in that adult thrips are only able to transmit TSWV if acquisition occurs in the first instar and early second larval thrips stages (Van De Wetering et al., 1996). The virus then replicates in the midgut and adjacent tissues, and eventually reaches the primary salivary glands (Ullman et al., 1992; Montero-Astúa et al., 2016; Rotenberg and Whitfield, 2018). After infection of the salivary glands, adults during feeding release the virus into viable plant cells along with the saliva. Hence, TSWV replicates in both the vector and the host plant, offering the opportunity for both direct and indirect (i.e. plant-mediated) effects on the vector. Studies of direct effects are rare (Belliure et al., 2005), but there are numerous reports of indirect effects of orthotospoviruses on thrips vector. For example, TSWV infection has been shown to increase vector fitness including, increased fecundity, development, population growth, and survival on infected plants relative to uninfected plants (Carter, 1939; Bautista et al., 1995; Maris et al., 2004; Belliure et al., 2005). Although most documented virus effects are positive, there are reports of negative (DeAngelis et al., 1993; Blua et al., 1994; Stumpf and Kennedy, 2005) and neutral (Wijkamp et al., 1996) effects of TSWV on thrips vectors as well. These reports suggest that TSWV infection can alter plant physiology to benefit vector fitness.

The literature on the molecular mechanisms underlying the tospovirus-thrips vector-plant interaction is emerging. Most studies have focused on defense-related signaling pathways as potential a mechanism in the model plants, *Arabidopsis thaliana* and *Nicotiana benthamiana*. Abe et al. (2012) demonstrated the role of antagonistic crosstalk between salicylic acid (SA) and jasmonic acid (JA)-responses in TSWV-thrips interaction using *Arabidopsis* mutants. The authors suggested that increased performance of thrips on TSWV-infected was caused by a reduction of the JA-regulated plant defense, which was suppressed by an increase in SA-regulated plant defense responses in virus-infected plants. More recently, Wu et al. (2019) showed that the TSWV non-structural protein, NSs protein suppressed biosynthesis of monoterpenes, which are known to repel *F. occidentalis* by directly interacting with MYCs, key regulators of the JA signaling pathway. To date, there have been four studies on transcriptional responses (microarray and RNA-Seq) to TSWV in host plants which report changes in plant immune defenses and metabolism in virus-infected plants (Catoni et al., 2009; Nachappa et al., 2013; Padmanabhan et al., 2019; Xu et al., 2020). However, these studies did not analyze global transcriptional changes in response to the thrips vector and the combination treatment of virus and vector.

The roles of the three major phytohormones, SA, JA and ethylene (ET) in defense responses to pathogens and insects is well-established (Walling, 2000; Stout et al., 2006; Howe and Jander, 2008; Bari and Jones, 2009; Pieterse et al., 2009; Verhage et al., 2010). However, research in the past decade demonstrates that plant defense responses is more than just SA and JA/ET pathways, with more coordination and integration of a range of hormones including, abscisic acid (ABA), auxins, brassinosteroids, cytokinins and gibberellins (De Bruyne et al., 2014; Shigenaga and Argueso, 2016; Yue et al., 2016; Berens et al., 2017; Yang et al., 2019). For instance, ABA can act as both a positive and negative regulator of disease resistance. ABA can suppress SA-mediated defenses and plant susceptibility to pathogens can increase following exogenous applications of ABA (Asselbergh et al., 2008). In certain instances, exogenous application of ABA can have the opposite effect on SA-mediated defenses resulting in increased resistance to pathogens (Ton and Mauch-Mani, 2004; Wiese et al., 2004; Melotto et al., 2006). The antagonism of SA-mediated defenses by ABA may be explained in part by the positive effect of ABA on JA biosynthesis (Adie et al., 2007). These phytohormones not only mediate immunity, but also growth and development (Pieterse et al., 2009; De Bruyne et al., 2014; Berens et al., 2017; Züst and Agrawal, 2017). For example, the classic growth hormone, auxin acts in an antagonistic manner with SA during plant defense, whereas auxin and JA signaling act synergistically. Moreover, some pathogens either produce auxin themselves or increase plant auxin biosynthesis to manipulate the plant’s defensive and development (Reviewed in (Kazan and Manners, 2009). The evolutionary conservation of the intricate network of phytohormone signaling pathways likely enables plants to effectively balance trade-offs between defense and growth (Züst and Agrawal, 2017).

In the current study, we sought to elucidate the plant-mediated mechanisms underlying the interaction between TSWV and its insect vector, *F. occidentalis* in the plant host, tomato, *Solanum lycopersicum* L. First, we performed replicated greenhouse and laboratory experiments to confirm that TSWV altered vector performance and behavior in ways that improved virus transmission. To characterize plant molecular mechanisms, microarray analysis was done in tomato plants that were systemically-infected with TSWV, infested with thrips, or both TSWV and thrips using the Affymetrix Tomato GeneChip®. The microarray represents myriad genes associated with primary metabolism, including chloroplast function, cell wall modification, protein synthesis, as well as defense- and stress-related genes associated with phytohormone signaling pathways (JA, SA, ET, ABA, and Auxin). In addition, we analyzed phytohormone levels and total free amino acid content in plants exposed to single and dual challengers. This is the first comprehensive analysis of plant-mediated mechanisms (genes and metabolites) that potentially improve the quality of TSWV-infected plants for its thrips vectors.

## MATERIALS AND METHODS

### Greenhouse experimental design and treatment structure

The greenhouse experiment consisted of four treatments: 1) TSWV infection alone; 2) thrip*s* infestation alone; 3) TSWV infection and thrips infestation, and 4) mock-inoculated, healthy controls. Each treatment was replicated three or four times (i.e., three or four plants) in a randomized complete block design with one plant per treatment per block. The experiment was conducted three times (i.e., three biological replication).

### TSWV and thrips sources

TSWV (isolate TSWV-MT2) was maintained by mechanical inoculations on caged tomato plants (*Solanum lycopersicum* cv Moneymaker) under greenhouse conditions as per (Rotenberg et al., 2009). A colony of thrips was maintained on green bean pods (*Phaseolus vulgaris*) as described by Bautista et al., (1995) at ambient room temperatures of 24 ± 1°C and a photoperiod of 16:8 h (L:D) light: dark cycle.

### Plant growth conditions, virus inoculations, and thrips release

Tomato plants (cv Moneymaker) were grown in 6-inch pots filled with Metro mix® potting soil and housed in one thrips-proof-screened (No Thrips Insect Screen®, BioQuip Products, Rancho Dominguez, CA) cage in a greenhouse room. Plants were fertilized once a week with Miracle Gro®-Water Soluble All Purpose Plant Food (24-8-16) NPK. The temperatures in the greenhouse ranged from 23-25 °C and the photoperiod was 16:8 h (L:D).

To generate TSWV-infected plants, three-week old plants were mechanically-inoculated with TSWV from infected plant tissue or mock-inoculated with buffer and healthy plant tissue. Virus inoculum was prepared by grinding two to three young symptomatic tomato leaves in ice-cold 5-10 ml of inoculation buffer (10 mM sodium sulfite and 5% wt/vol celite) using a pre-chilled mortar and pestle. Inoculum was applied and dragged lightly with a cotton swab over the surface of all fully-expanded leaves on the plant. Approximately two weeks after inoculation, plants were visually inspected for TSWV symptoms, i.e., stunting and deformation, chlorotic ring spots, mosaic patterns, and leaf-bronzing, and the most uniform group of symptomatic plants were chosen for each experiment.

Individual 5-week-old symptomatic and mock-inoculated control plants were moved to single-plant cages constructed from 19-liter cardboard ice cream buckets (38 cm tall × 26 cm diameter) with four 14 × 27 cm apertures cut into the side walls. The four apertures were covered with No-Thrips Insect Screen™ (GreenTek, Inc., Edgerton, WI, USA) and sealed with silicone. The top of the container was covered by thrips-proof-screen secured by rubber bands. These single-plant cages prevented cross-contamination between treatments. Cohorts of 40 adult females from the laboratory colony, 7-d post eclosion were transferred into 1.5 ml microcentrifuge tubes and then allowed to escape in a synchronous manner from one tube placed at the base of each plant.

### Thrips feeding damage, performance and settling assays

Thrips feeding damage was quantified using a visual rating scale according to McKinney (1923). All leaves from each plant were evaluated visually and grouped into six classes with values of 0, 1, 3, 5, 7 and 9, which corresponded to percentages of total leaf area damaged by thrips feeding of 0, < 10%, 10–25%, 25–50%, 50–75% and > 75%, respectively. The leaf damage index values were calculated according to the following formula: LFDI (%) = 100 × (Σ class value × corresponding leaf numbers)/(total numbers of leaves × 9). In addition, the number of thrips feeding lesions per plant were also counted.

Thrips performance on virus-infected plants or mock-inoculated plants was defined as the number of offspring that emerged from leaf tissue at 7-d post release of adult thrips. For each biological replicate, 2-sample rank tests (i.e., Mann-Whitney) using Minitab v.14 (Minitab, Inc., State College, PA) was performed to determine if the median count of offspring obtained from TSWV-infected plants was similar to that obtained from healthy, mock-inoculated plants. Data from all three experiments were pooled together and a two-way analysis of variance in Minitab using a GLM with treatment and biological replicate as fixed effects.

To determine adult thrips settling preference, assays using detached-leaflets were conducted as per (Nachappa et al., 2013). Same-age leaflets were taken from mock-inoculated plants and plants with TSWV infection alone at the termination of each greenhouse experiment (*i.e.*, 6-week-old plants). Leaflet pairings consisted of choice and non-choice tests using pairs of leaflets in 15-cm diameter Petri-dishes with the lids of each dish fitted with thrips-proof screen to allow ventilation. Each Petri dish contained 15 ml of 1.5% water agar onto which were placed the adaxial surfaces of each paired leaflet. The agar prevented desiccation of leaflets while allowing thrips to move across the surface and choose leaflets. Ten adult female thrips were placed with a small paint brush in the center of the Petri dish, equidistant from each leaflet, and lids were sealed with Parafilm. The assay was conducted under laboratory conditions with a 16:8 h photoperiod and ambient temperatures of 24-25°C. Adult thrips preference (choice of virus infected vs. mock controls) was determined by counting the number of thrips present on each of the paired leaflet every hour for the first six hours, and then at 12, 24, 36, 48, and 72 h after the release. For each biological replicate, we performed 1-Sample Wilcoxon sign rank tests using Minitab to determine if the median paired difference in accumulation on virus-infected vs. non-infected healthy tissue was significantly greater than zero at each time-point.

### Leaf tissue sampling for gene expression, phytohormone and amino acid analyses

Preliminary observations revealed that thrips preferred to feed on older (basal) leaves, so leaflets were harvested from 10^th^ youngest leaf, counting down from the top to the base of the plant. From each of the three experiments (biological replications), two same-age, paired leaflets (on each side of leaf rachis) were harvested. One leaflet was processed for gene expression analysis (i.e., microarray hybridizations and RT-qPCR analysis) and the other was processed to determine phytohormone contents. Leaflet samples were flash-frozen in liquid nitrogen and stored at −80°C. To obtain enough leaf tissue (approximately 5 g) required for determination of total free amino acid content, a third leaf immediately basal to the leaflets chosen for microarray and phytohormone analyses was simultaneously harvested and freeze-dried.

### RNA isolation, amplification, labeling and generation of GeneChip data

Total RNA isolation and cDNA synthesis was performed as per Nachappa et al. (2013). Briefly, RNA was isolated from frozen leaflet samples (100 mg of tissue) using the Qiagen RNeasy Plant Mini Kit (Qiagen, Valencia, CA, U.S.A) following manufacturer’s protocol. RNA quantity was determined by NanoDrop spectrophotometer (NanoDrop Technologies, Wilmington, DE, U.S.A.) and quality was assessed with the 2100 Bioanalyzer using Nanochip technology (Agilent Technologies, Inc., Palo Alto, CA U.S.A.). Total RNA was pooled for each treatment (3-4 experimental replicates, i.e., plants) within a biological replication. Pooled RNA samples were subjected to cRNA synthesis, labeling, and hybridization to Affymetrix Tomato Genome Arrays (GeneChip®) at Kansas State University, Integrated Genomics Center. Each GeneChip® contains more than 10,000 probe sets for over 9,200 genes with each gene being represented by at least one probe set containing 25-mer oligonucleotides. Hybridization intensities of scanned microarrays for each of the three biological replicates were generated with Gene Chip Operating System, GCOS (Affymetrix Inc.). Global scaling was applied for each GeneChip® to adjust the Target Intensity (TGT) Value to an arbitrary target of 500 so that hybridization intensity of all chips was equivalent. In addition, expressed genes were identified by GCOS, using a detection algorithm and assigned a present, marginal, or absent call for genes represented by each probe set on the array (GeneChip Expression Analysis Technical Manual). Microarray data files (.CEL) were analyzed using GeneSpring 10.1 (Silicon Genetics, Redwood, CA, U.S.A.) and normalized using RMA (Robust Multichip Average) algorithm. Differentially-expressed genes were selected using 2 criteria: i) an expression ratio of at least ± 2-fold change and ii) *P* ≤ 0.05 in ANOVA tests comparing log2 (normalized hybridization intensity) of treatment to the mock control. Differentially-expressed genes were assigned functional annotations with Blast2GO software (Conesa and Götz, 2008)) and classified by GO-biological process, cellular component and molecular function using default parameters and an *E*-value cut-off of 10^-5^. The microarray experiment design details and raw microarray data is available at ArrayExpress under the accession number E-MTAB-9294.

### Hierarchical cluster analysis

We used the heatmap () base function in R to create a heatmap of a sub-set of 369 differentially expressed genes selected for their membership in candidate pathways (defense, phytohormones, photosynthesis, cell wall organization, and protein metabolism). The colors range from red to green, showing the expression of genes in each treatment group ranging from highly similar (green) to less similar (red). The heatmap was plotted from the adjacency matrix of the network, using the R heatmap () function. The Pearson correlation coefficients depicts correlation between genes and treatments based on the average fold change values from 3 replicates for each treatment.

### Phytohormone signaling pathway analysis

We analyzed changes in tomato phytohormone pathways for SA, JA, ET, ABA and AUX-responses associated with each treatment using the pathway analysis method developed by Studham and MacIntosh (2012). This method provides a broad indication (pathway score) of relative upregulation or downregulation of individual phytohormone pathways based on the magnitude (fold change), direction of change (up or down), and significance of differentially-expressed genes (compared to mock treatment) associated with each pathway for each treatment. Pathway scores for each phytohormone pathway can be compared to provide a comprehensive picture of treatment-associated, host plant responses and to identify dominant pathways affected by treatment. To perform the pathway analysis, annotations for probe sets in the Tomato GeneChip® were obtained manually based on literature and from Blast2GO biological processes (https://www.blast2go.com) (Conesa and Götz, 2008). Additional annotations for probe sets that were not assigned a function were obtained by identifying the EC numbers using the annot8r program (http://www.nematodes.org/bioinformatics/annot8r/). The EC numbers were used to identify pathway membership by querying UniProt (https://www.uniprot.org/) and KEGG (Kanehisa et al., 2011). The genes associated with different functional categories have different roles such as biosynthesis, response, regulation, and others. Each role was assigned different weights indicating its relative importance in determining the induction or suppression of a hormone/protein (Studham and MacIntosh, 2012). A cumulative pathway score was then calculated for the subset of genes that exhibited microarray fold changes ≥ 1.0 and p-values ≤ 0.05 for each phytohormone pathway.

### Co-expression network analysis

Gene co-expression network analysis is used to describe the correlation patterns among genes across microarray and other multidimensional expression data sets (e.g. RNAseq). The weighted gene correlation network analysis method (WGCNA, Langfelder and Horvath, 2008) can be used to identify clusters (modules) of highly correlated genes, and each module provides information about the pairwise relationships (correlations) between genes. In addition, community analysis of gene co-expression networks can identify communities (clusters, modules) of nodes. The nodes in a particular community have a higher likelihood of connecting to each other than to nodes from other communities (Barabási, 2016), and these relationships can provide additional information about groups of genes that are likely to be expressed together under particular experimental conditions. The WGCNA R package [https://cran.r-project.org/web/packages/WGCNA/index.html; Langfelder and Horvath, 2008] was used to analyze a sub-set of differentially expressed genes in candidate functional categories of interest (*i.e*, photosynthesis, plant hormone-related (ABA, AUX, ET, JA, SA), protein metabolism and turnover, cell wall and defense functions) across four treatments (mock-infected, TSWV, thrips and TSWV+thrips). The sample code in the WGCNA tutorial (Langfelder and Horvath, 2008) (https://horvath.genetics.ucla.edu/html/CoexpressionNetwork/Rpackages/WGCNA/Tutorials/index.html) was adapted to perform WGCNA on our expression data set. The input data to WGCNA consisted of averaged three replicates for each of the four treatments. Code from the first tutorial on WGCNA network analysis (https://horvath.genetics.ucla.edu/html/CoexpressionNetwork/Rpackages/WGCNA/Tutorials/FemaleLiver-02-networkConstr-auto.R) was adapted and used to construct the correlation network. The nodes and edges of the WGCNA network were exported to files using the export network to Cytoscape (Shannon et al., 2003) function in the WGCNA package. We tried several network visualization tools, including the popular Cytoscape and Gephi (Bastian et al., 2009). However, we needed a bit more flexibility than these tools provided, so the network figure was created using the R igraph package (https://igraph.org/r/). Different community detection methods were applied to the network data to determine the optimal method and number of community modules, and the leading eigen method was selected (Newman, 2006). As our goal was to visualize the network nodes clearly, we tried most of the igraph network layouts and compared the resulting network images. The ‘layout_with_fr’ igraph layout was chosen for the figure because it produced the clearest visualization of the network. This layout uses the force-directed layout algorithm of Fruchterman and Reingold to place the network vertices on a plane (Fruchterman and Reingold, 1991). The resulting network included five communities of nodes, each community representing a functional category (see list above).

### Confirmation of microarray data using reverse transcription quantitative –PCR (RT-qPCR)

We targeted six genes associated with the SA, JA, and antiviral small-RNA-mediated gene silencing pathways. These genes were BGL2 and NPR1 (SA pathway) and OPR3, AOS and CI (JA pathway), and RNA-directed RNA polymerase 1 (RDR1). We chose elongation factor 1-alpha (*le*EF1) as the internal reference gene for normalization because expression of this gene was found to be invariant with TSWV infection or herbivore challenge in the microarray experiments and had been previously shown to be stably-expressed in Moneymaker tomato systemically-infected with tobacco rattle virus (Rotenberg et al., 2006). Target and leEF1 primer pair sequences, their corresponding melting temperatures, and real-time PCR efficiencies are indicated in Table 1. The normalized abundance of TSWV nucleocapsid (N) RNA compared to leEF1 was also determined to estimate virus titer in leaf tissue using primers tested previously (Rotenberg et al., 2009). We selected one plant per treatment (mock included) per biological replicate (*i.e*., 24 RNA samples in total) of the greenhouse experiment that represented the average fecundity and/or TSWV symptom severity for a given treatment. Subsamples of total RNA isolated from leaflet tissue used in the microarray hybridization experiment were treated with DNase using the rigorous DNA removal procedure of the Turbo DNA-free kit (Applied Biosystems Inc, Carlsbad, U.S.A) and cDNA was synthesized from 1 µg DNA-free RNA using the iSCRIPT cDNA synthesis kit (BioRad, Hercules, CA, U.S.A). Real-time PCR master mixes were prepared using iQ sybr Green Mix (BioRad) according to manufacturer’s specifications and final reaction (20 µl) concentrations of 200 nM of each primer. Reactions were performed in duplicate using the iCycler iQ Thermal Cycler with a 96 x 0.2 ml reaction module and iCycler iQ software (Bio-Rad).

**Table 1.**
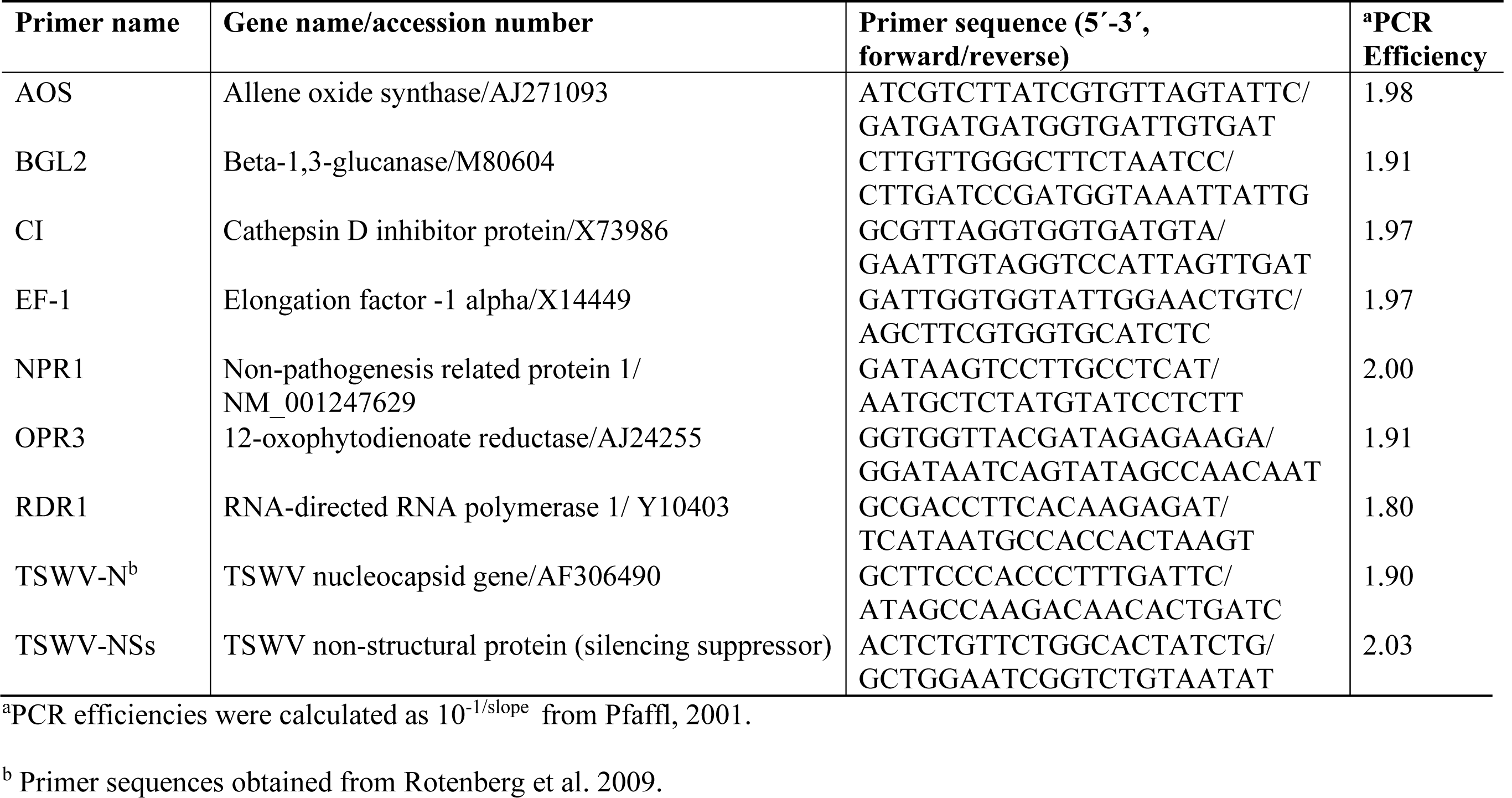
Real-time quantitative reverse transcriptase-PCR primer pair sequences and corresponding PCR efficiencies.

The relative abundance of target RNA was determined for each virus and herbivore treatment compared to the mock control. The relative expression ratio (RER) equation (Pfaffl, 2001) was calculated as follows: RER = E_target_^ΔCttarget[control-treatment]^/ E_ref_ ^ΔCtref[control-treatment]^ where E refers to the PCR primer efficiency for target or internal reference (ref; leEF1) genes and ΔCt is the difference in Ct-values (i.e., threshold cycle values automatically calculated by the IQ software) between a treatment and mock control. The average Ct value (n = 3) obtained for the mock control for each target and reference gene was determined and used in the RER calculations. To estimate virus titer, the normalized abundance of TSWV N or NSs RNA (genomic and transcript RNA) was calculated using the Pfaffl inverse equation (Pfaffl, 2001): E_ref_^Ctref^/E_N_^CtN^ as described previously (Rotenberg et al., 2009).

### Phytohormone analysis

Frozen tomato leaflet samples from each biological replicate were sent to The Donald Danforth Proteomics & Mass Spectrometry Facility, St. Louis, MO for chemical extraction and quantification of SA, JA, ABA, jasmonyl-isoleucine (JA-IL), and 12-oxo-phytodienoic acid (OPDA) using liquid chromatography–electrospray tandem mass spectrometry as per (Pan et al., 2008). Analysis of variance was performed on log_10_-transformed phytohormone contents (ng analyte g fresh weight^-1^) in Minitab using a GLM that included treatment and biological replicate as two fixed factors and their interaction term as the random factor. The analyses revealed no apparent main effect or treatment interaction for any of the phytohormones measured due to biological replicate; therefore, data was combined for the three biological replicates and one-way ANOVA was performed to determine the main effect of treatment on phytohormone concentration. Pairwise treatment comparisons were performed using Tukey’s family error rate (*P* ≤ 0.05).

### Total free amino acid analyses

Frozen tomato leaf samples obtained from each experiment were sent to the Ruminant Nutrition Laboratory at Kansas State University for extraction and quantification of total free amino acids. Total free amino acid content was analyzed using a modified protocol described (El Fahaam et al., 1990). Briefly, whole leaves were harvested from tomato plants and freeze-dried in an oven/ desiccator at 80°C for 2-3 days or until no change in weight was recorded. Approximately 0.1g of dry tissue was extracted with 10 ml of 70% hot ethanol and centrifuged at 2500 *g* for 5 min. The dry residue was dissolved in 2.5 ml of 0.1 N HCl and kept at −20°C until assayed. Colorimetric procedures were adapted to Technicon Autoanalyzer II for simultaneous determination of total free amino acid content based on an internal standard (leucine) in plant tissue samples (modified from (Broderick and Kang, 1980). Total free amino acid content data were analyzed using a similar statistical model as the phytohormone analysis; however, biological replicate had a significant main or treatment interaction effect. Therefore, data from biological replicates were interpreted separately.

## RESULTS

### Performance and settling behavior of thrips

The effect of TSWV infection of tomato plants on thrips performance (number of offspring) was evaluated seven-days after adult females were released onto individual plants. Our data revealed significant differences in the number of offspring (first and second instar larvae) produced on virus-infected plants compared to mock-inoculated plants (F= 11.78, df=1, *P* = 0.003) (**Fig. 1A**). There was no impact of time or biological replication (F= 1.73, df=2, *P* = 0.21) and the interaction between number of offspring and time (F= 0.03, df=2, *P* = 0.97) on thrips population. On average, there were 2-fold more thrips offspring on virus-infected plants compared to mock-inoculated plants (Mean ± SE: 25.75 ± 2.02 and 13.50 ± 2.67, respectively). Leaf damage caused by thrips was quantified and no apparent differences were detected between thrips on healthy or virus-infected plants (leaf damage index: *P*=0.34; number of lesions: *P*=0.48, Kruskal-Wallis test). In Petri dish assays, thrips adults were given a choice between TSWV-infected and mock-inoculated leaflets harvested from the greenhouse experiments. By 3 to 4-h post-release, there were significantly more thrips adults (Z= 10.0, *P=*0.03 and Z= 3.0, *P*=0.02, respectively) associated with TSWV-infected leaflets compared to mock-inoculated leaflets (**Fig. 1B**). This trend persisted over the course of the 72-h experiment. There were no apparent differences between the numbers of thrips observed between leaflets in the non-choice situation (*P* > 0.2 for all time-points, data not shown).

**Fig. 1.**
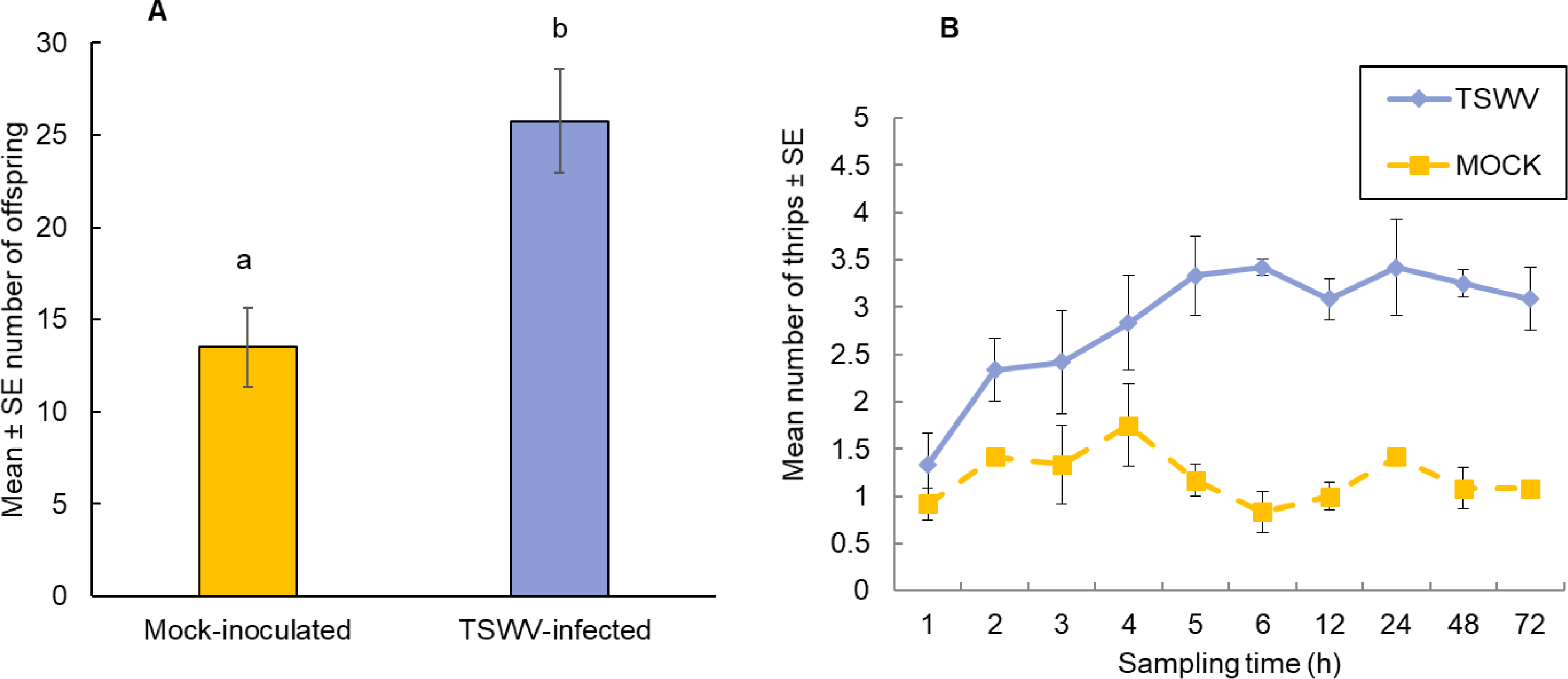
Performance and settling preference of thrips, *F. occidentalis*, on tomato plants that were infected with TSWV or mock-inoculated. A) Mean number of thrips offspring (first and second instars) on tomato plants one-week after adult female release. Each bar represents the average of n=4 plants per experiment or biological replicate, and B) Number of adult female thrips recovered on TSWV-infected and mock-inoculated leaflets in detached Petri dish assays. Leaflet pairs were obtained from same-aged TSWV-infected and mock-inoculated tomato plants from the corresponding greenhouse experiment. Values at each time point represent the average of n=3-4 leaflets originating from a different greenhouse experiment or biological replicate. Plants were inoculated with TSWV by leaf-rub inoculation two weeks prior to insect release. Different letters indicate significant differences between treatments at *P*<0.05.

### Single and combined effects of TSWV and thrips on tomato global gene expression profiles

Tomato microarray hybridizations were performed to describe and quantify single and combined effects of virus and vector feeding on transcription-level expression. Collectively, of the 10,209 probe sets (i.e., 9,200 unique coding sequences) represented on the Tomato GeneChip^®^, 1,722 sequences were differentially-expressed in plants challenged by the various treatments compared to mock-inoculated, healthy plants (*P* < 0.05, regardless of magnitude of fold change). Of these sequences, 307, 171, and 424 genes were significantly expressed by at least 2-fold and *P*≤ 0.05 in TSWV, thrips, and TSWV+thrips challenged plants compared to the mock-inoculated controls (Supplementary tables 1, 2 and 3, respectively). Venn diagrams depicting the number of unique and shared genes that were differentially-expressed among treatments revealed several patterns (**Fig. 2A and B**). First, only a small proportion of genes were unique to individual challengers. Second, systemic infection of tomato plants by TSWV contributed the most to global gene expression changes in both the positive (up-regulation) and negative (down-regulation) direction as compared to changes induced by thrips feeding. Third, the majority of genes differentially-expressed in response to thrips alone were down-regulated (64%). Fourth, the combined effect of TSWV and thrips resulted in a greater proportion of down-regulated genes that were unique to dual challenger (i.e., 32% down vs. 16% up) (**Fig 2A and B**).

**Fig. 2.**
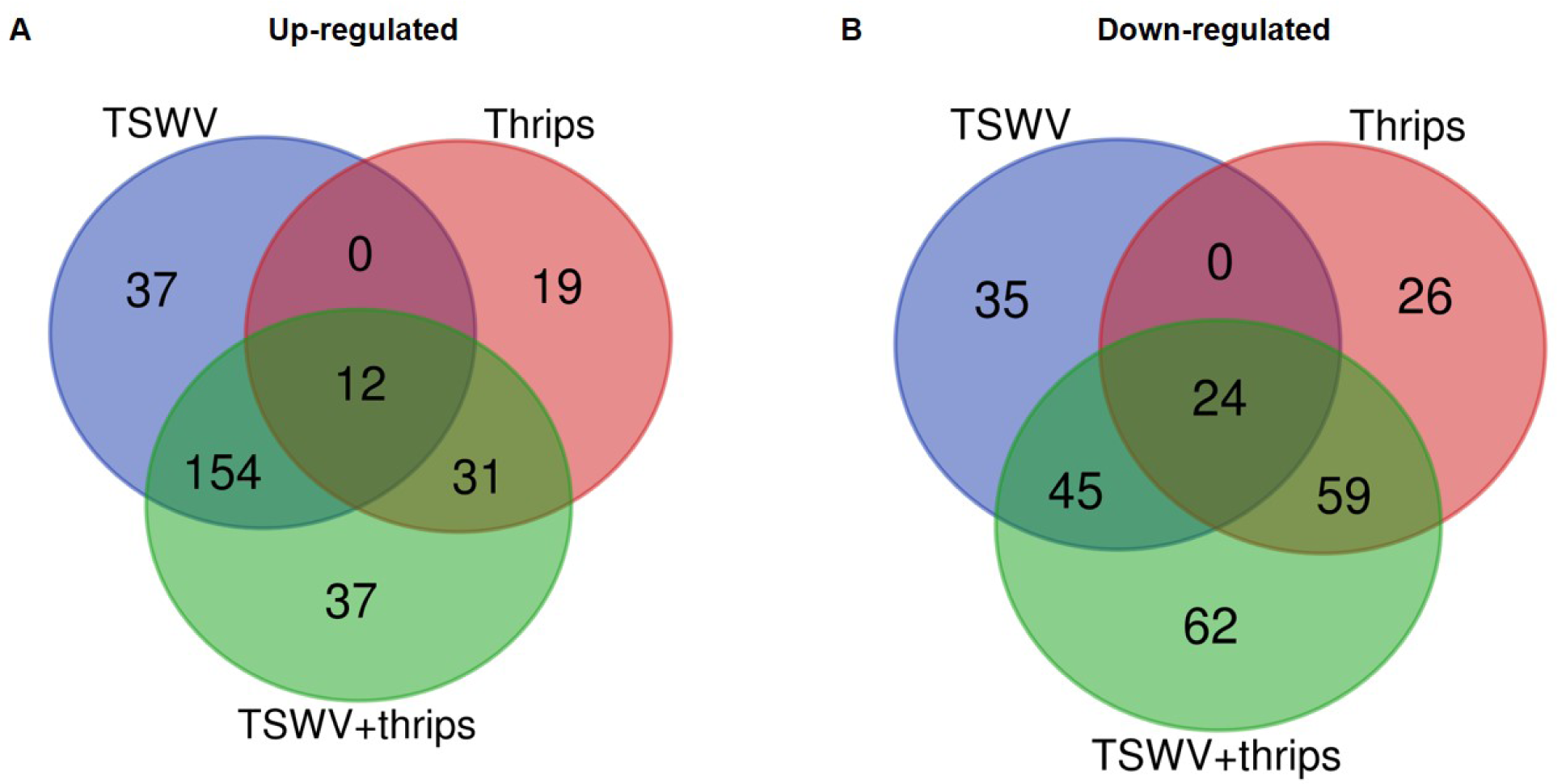
Venn diagrams depicting number of unique and shared differentially-expressed genes. Data were obtained from differentially-expressed genes in tomato plants systemically-infected with TSWV and/or infested with *F. occidentalis*. Numbers outside of circles indicate the total number of differentially-expressed genes for a particular treatment.

**Table 2.** Relative activity of phytohormones in the TSWV- *Frankliniella occidentalis* interaction. Phytohormone pathway scores indicate induction (positive), suppression (negative), or insignificant effect (less than 1) of signaling pathways in response to virus infection, thrips infestation and the combination treatment.

| **Treatment** | **Salicylic acid** | **Jasmonic acid** | **Ethylene** | **Abscisic acid** | **Auxin** |
| --- | --- | --- | --- | --- | --- |
| **TSWV** | 62.25 | -58.28 | 0.24 | 27.05 | -77.83 |
| **Thrips** | -38.07 | 40.02 | 40.01 | -16.8 | 6.57 |
| **TSWV + thrips** | 37.62 | 56.99 | 0.64 | 76.36 | -48.63 |

**Table 3.** Reverse transcription quantitative-PCR (RT-qPCR) validation of differential genes. Relative transcript abundance in tomato plants systemically-infected with TSWV and/or infested with thrips. Values represent mean of three biological replicates. Different letters indicate statistical difference between treatments at *P* ≤ 0.05.

| Gene title | Probe set ID | Log<sub>2</sub>(Fold change intensity) |  |  | Log<sub>2</sub>(Relative expression ratio) |  |  |
| --- | --- | --- | --- | --- | --- | --- | --- |
|  |  | TSWV | Thrips | TSWV + thrips | TSWV | Thrips | TSWV + thrips |
| **<u>SA pathway</u>** |  |  |  |  |  |  |  |
| Beta 1,3 glucanase | Les.3673.1.S1_at | 3.6a | -0.9a | 3.4a | 9.2a | 3.3b | 7.3a |
| Non-expressor of pathogenesis-related 1 | Les.5940.1.S1_at | 0.7a | -0.4b | 0.6a | 2.3a | -0.2b | 1.0a |
| **<u>JA pathway</u>** |  |  |  |  |  |  |  |
| Allene oxide synthase | Les.13.1.S1_at | -1.4a | 0.6a | -0.04a | 1.0a | 0.3a | 1.5a |
| Cathepsin D inhibitor protein | Les.3740.1.S1_at | -2.4b | 0.9a | 2.4a | -2.8b | 4.2ab | 5.1a |
| 12-oxophytodienoate reductase | Les.22.1.S1_at | 0.8a | 0.2a | 0.8a | 2.4a | -0.2b | 1.4ab |
| **<u>RNAi pathway</u>** |  |  |  |  |  |  |  |
| RNA-directed RNA polymerase 1 | Les.61.1.S1_at | 1.1a | -0.04b | 1.1a | 3.8a | 1.2b | 3.5a |

The differentially-expressed genes were functionally-classified by Gene Ontology (GO) terms into biological process, cellular component and molecular function with relevance to plant health and responses. Overall, a greater proportion of genes responded to TSWV infection alone and combined treatment compared to thrips feeding alone in all three GO categories (**Fig. 3A, B and C**). The GO-biological process most represented by the differential expression was in response to stimulus, including biotic and abiotic stimulus (**Fig. 3A**). Within this category, a large percentage of genes were up-regulated in response to virus infection alone (74%) and the combination treatment (64%), whereas thrips feeding down-regulated a larger percentage of stress-related genes (63%). The second largest category was photosynthesis, and a majority of photosynthesis-associated genes (90%) were down-regulated in all three treatments. This trend is also reflected in GO cellular component category where chloroplast-associated genes were over-represented, and a large percentage (95%) were down-regulated across all three treatments (**Fig. 3B**). Another category of potential interest was cell wall organization, which was overrepresented in GO-cellular component and GO-biological process as well (**Fig. 3A and B**). Virus infection down-regulated 25% of cell wall related genes, but some genes were also up-regulated (14%). With regards to GO-molecular function, ATP binding genes were over-expressed, which agrees with the GO-CC category where cell membrane-associated genes were overrepresented (**Fig. 3C**). There was no trend in ATP binding genes in response to virus infection, in contrast, thrips feeding resulted in down-regulation of ATP binding genes (100%). The protein binding category was overrepresented, with virus infection in the single and combined treatment resulting in induction of protein binding genes (69% and 87%, respectively), whereas thrips feeding resulted in suppression of the genes (88%).

**Fig. 3.**
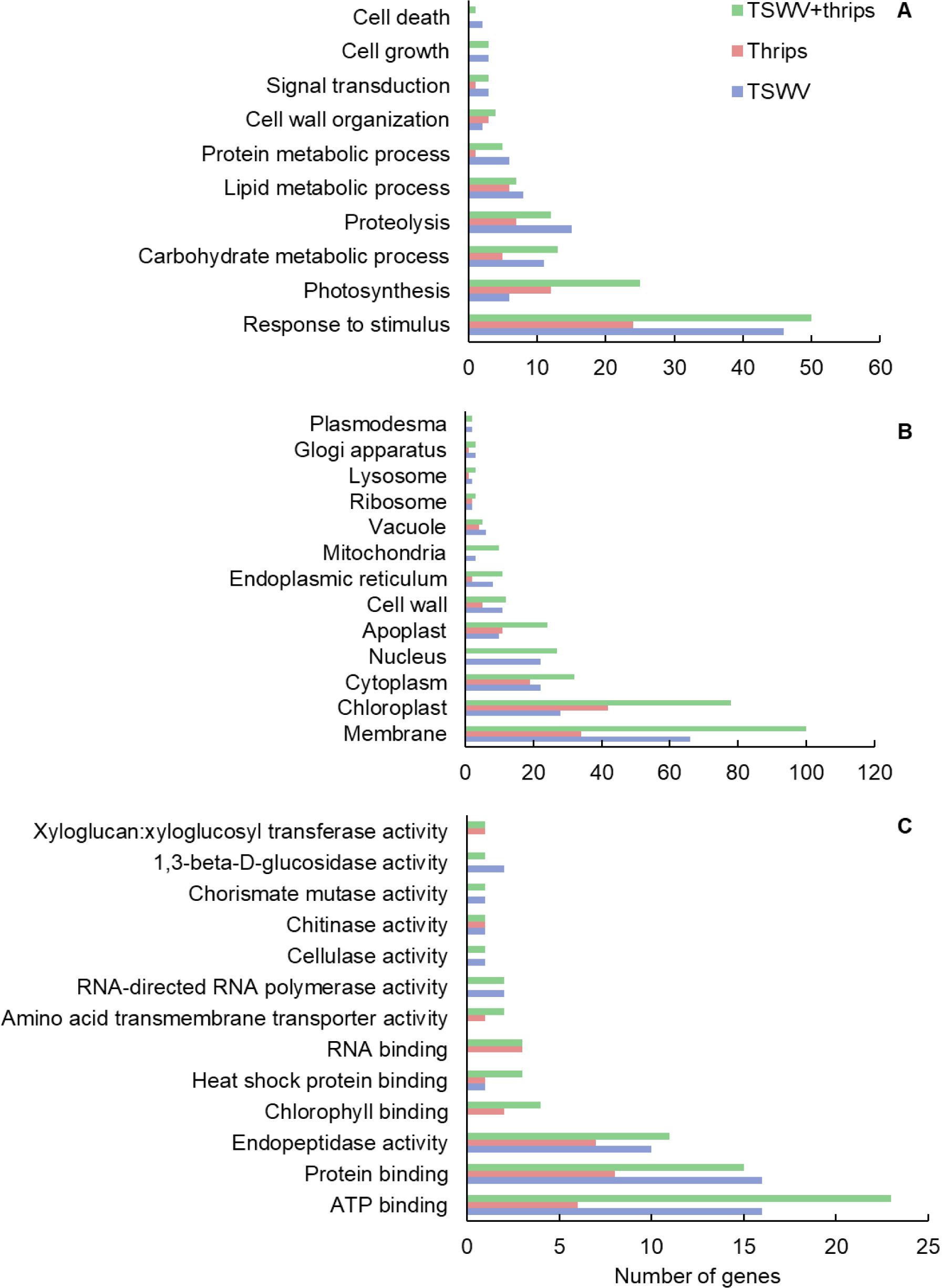
Gene ontology (GO) terms for differentially-expressed genes. Distribution of differentially-expressed genes in tomato plants systemically-infected with TSWV and/or infested with *F. occidentalis*. (A) Biological Process, (B) Cellular Component, and (C) Molecular Function.

### Single and combined effects of TSWV and thrips on defense-related and primary metabolic processes inferred from gene expression

#### Defense signaling pathways

We examined the microarray hybridization data for possible interaction between SA and JA pathways. The magnitude and direction of differential expression of all known phytohormone genes in the Tomato GeneChip are shown in **Supplementary Table 4**. TSWV infection alone significantly up-regulated the majority (59%) of signature SA-responsive genes compared to mock-inoculated plants (**Fig. 4A**). These include genes involved both upstream and downstream of SA synthesis such as chorismate mutase and pathogenesis-related proteins, respectively. Genes encoding the proteins NPR1 and NPR3 that have been shown to interact with SA were also up-regulated in virus-infected plants. We observed suppression of JA-genes in response to virus treatment; with 29% suppression and 18% up-regulation in virus-infected plants (**Fig. 4B**). This suggests that the strength of the SA-induced suppression of JA-genes was not as widespread. Moreover, most of the down-regulated JA genes belonged to the protease inhibitor category specifically, wound-induced proteinase inhibitors. A large percentage (60%) of ET-associated genes including those involved in ET synthesis such as 1-aminocyclopropane-1-carboxylate oxidase and ET signaling such as ethylene-responsive transcription factor 5 were up-regulated in TSWV-infected plants (**Supplementary Table 4**). Feeding by thrips alone resulted in up-regulation of JA-related genes (70%), but no major impact was observed on ET genes (**Fig. 4B**, **Supplementary Table 4**). Interestingly, genes in all three signaling pathways were significantly up-regulated in TSWV and thrips dual treatment **(Supplementary Table 4).** In addition to signaling pathways, genes involved in general stress responses such as heat shock proteins, GSTs, and reactive oxygen species (ROS) were differentially-regulated in TSWV infected tomato, and in most cases up-regulation occurred regardless of the presence of thrips **(Supplementary Table 4)**. We also found that transcription factors such as WRKYs, Myb family, bZIP family and Mitogen-activated kinase 6 that initiate defense responses to be up-regulated in response to virus infection.

**Fig. 4.**
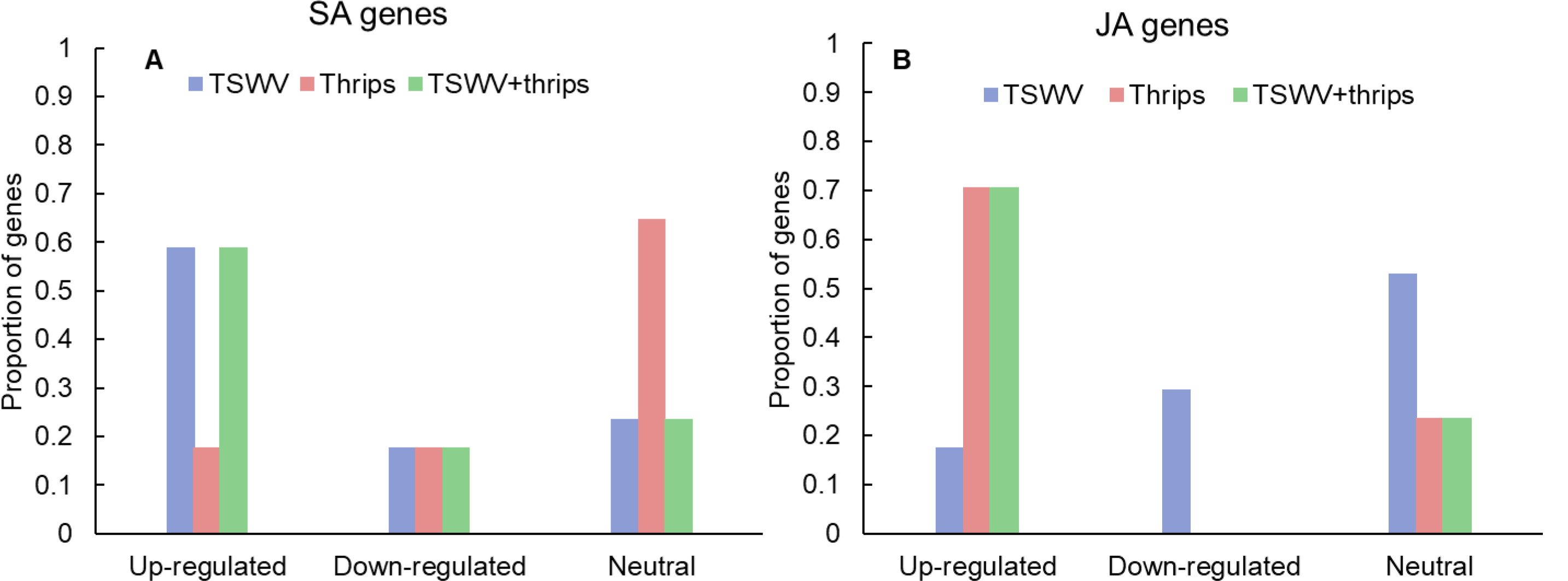
Proportion of differentially-expressed genes in the A) salicylic acid (SA) and B) jasmonic acid (JA) pathway in response to TSWV infection alone, *Frankliniella occidentalis* alone, and the combined treatment. n = 17 possible SA genes, n = 17 possible JA genes represented on the array. The list of SA and JA genes and the average normalized hybridization values is provided in Supplementary Table 4.

We analyzed other phytohormones that are known interact with SA and JA/ET, namely ABA and auxin and that were differentially-expressed in our microarray analysis. Virus infection alone did not significantly impact ABA-related genes (66% showed similar expression to mock) and 33% ABA-related genes were down-regulated (**Supplementary Table 4**). Thrips alone and the dual treatment did not differ significantly in the expression from mock (83% and 60% similarity, respectively) (**Supplementary Table 4**). With regards to auxin, TSWV infection alone and in combination with thrips down-regulated 40% and 60% of auxin genes, whereas thrips activity had similar percentage of up-regulated and down-regulated genes (40% for both) (**Supplementary Table 4**).

#### Photosynthesis-related processes

Genes involved in photosynthesis were largely down-regulated in all three treatments (**Fig. 5A**, **Supplementary Table 5**). Virus infection, both alone and in combination with thrips feeding, had a greater impact on photosynthesis-related genes with 54% and 70% of the genes being suppressed compared to mock-inoculated plants, respectively (**Supplementary Table 5**). These include ribulose bisphosphate carboxylase, ATP binding protein, chaperone, and chlorophyll a/b-binding proteins. Plants challenged with thrips also repressed 37% of photosynthesis-related genes compared to the control.

**Fig. 5.**
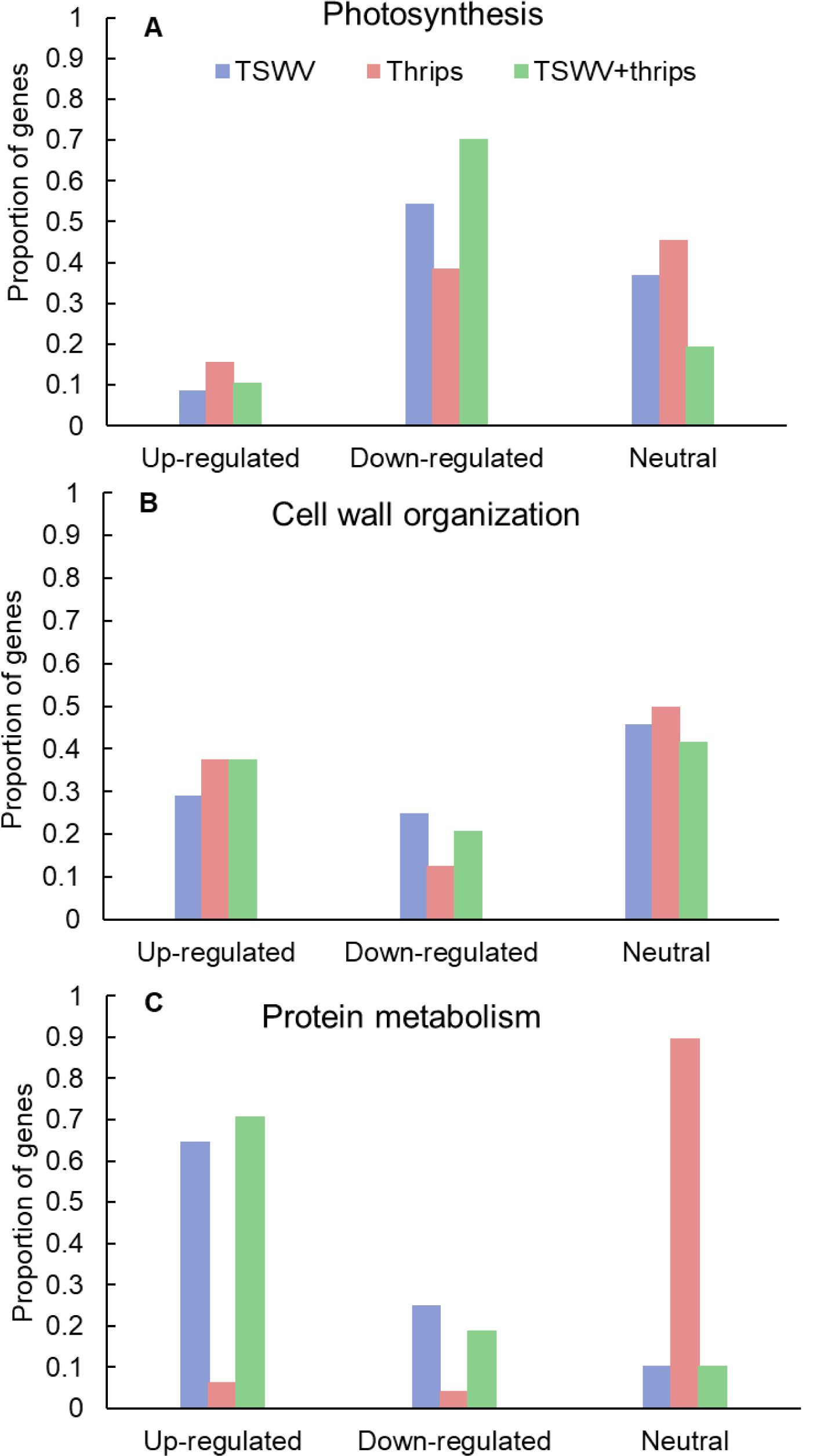
Proportion of differentially-expressed genes involved in A) photosynthesis and B) cell wall organization and C) protein metabolism in response to TSWV infection alone, *Frankliniella occidentalis* alone, and the combined treatment. n = 57 possible photosynthesis genes, n = 24 possible cell wall related genes and n=48 possible protein metabolism genes represented on the array. The list of these genes and the average normalized hybridization values is provided in Supplementary Table 5.

#### Cell wall organization

Virus infection altered gene expression in both upward and downward direction. Among the up-regulated genes (29%), cell wall degradation genes such as expansin, some cellulases and beta-1, 4-glucanases that are also involved in the SA-pathway (**Fig. 5B; Supplementary Table 5**). In contrast, cell wall-related genes that were significantly down-regulated (25%) were cell wall modification enzymes as xyloglucan endotransglycosylase/hydrolases and pectinesterase. In contrast, thrips feeding alone and combined treatment resulted in up-regulation of a large percentage of cell wall genes (37% for both; **Supplementary Table 5**). This perturbation of cell wall gene expression shows that thrips feeding has a major impact on cell wall genes even in the absence of virus infection in host tissues.

#### Protein synthesis and degradation

A majority of genes involved in protein synthesis such as constituents of 40S, 50S and 60S ribosomal subunits and genes associated with protein degradation such as those that encode 26S proteasome and involved in ubiquitination were induced in response to TSWV and the combination treatment (64% and 70%, respectively; **Fig. 5C** and **Supplementary Table 5**). The presence of thrips alone did not have an impact on expression of protein metabolism genes compared to mock-inoculated plants (**Fig. 5C**).

### Hierarchical cluster analysis

Focusing on the 369 DE genes associated with pathways of interest, the cluster analysis revealed more similar global gene expression patterns between TSWV infection alone and TSWV + thrips, and likewise, mock-inoculated and thrips alone treatments produced similar patterns (**Fig. 6; Supplementary Table 6**). Genes that were consistently up-regulated by virus alone or the dual treatment were protein metabolism genes such as E3 ubiquitin-ligase, 40S, 50S, and 60S ribosomal subunits (cluster 1, 5 and 6), but also involved genes with broad function in defense and phytohormone pathways, such as NPR1, PR-5 and EDS1 (**Supplementary Table 6**). In contrast, virus alone and the dual treatment consistently down-regulated photosynthesis and cell wall functional categories such as chlorophyll a-b binding proteins, rhodanese-like domain-containing proteins and xyloglucan endotransglycosylase/hydrolases, respectively (cluster 2, 3 and 4). Interestingly, TSWV alone and the mock treatment showed similar trends in expression (down-regulation) of the genes in cluster 7, which was predominantly comprised of JA and ET pathway genes (**Fig. 6**). Moreover, in this cluster of genes, thrips activity alone resulted in an expression pattern (up-regulation) that was more similar to that of the dual treatment, which included defense and phytohormone related genes. Thrips activity largely down-regulated genes related to protein metabolism, photosynthesis and cell wall functional categories (cluster 1, 4 and 5). These results provide insights into correlation between biochemical pathways, including photosynthesis, protein metabolism, cell wall biogenesis and defense during single and dual attack by TSWV and the thrips vector.

**Fig. 6.**
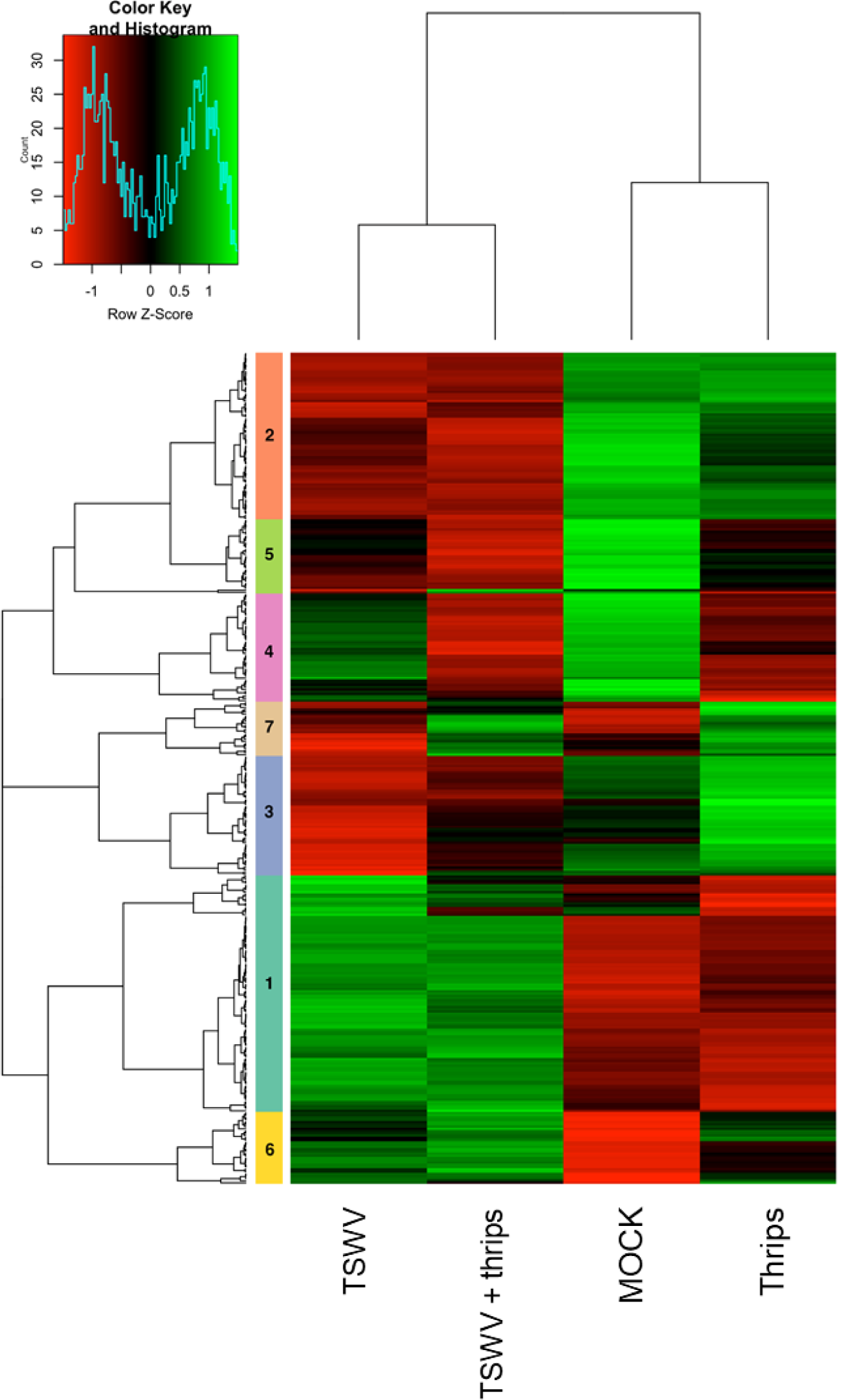
Hierarchical clustering of differentially-expressed genes in candidate pathways including defense and phytohormones, photosynthesis, cell wall organization, and protein metabolism. Heatmap showing the correlations of genes and treatments based on the averaged normalized hybridization data from 3 replicates per treatment. The row clusters indicate genes that are correlated in terms of their expression and treatment from green (positive correlation) to red (negative correlation). The list of genes in each cluster is provided in Supplementary Table 6.

### Pathway analysis

Pathway analysis was performed on the microarray data to determine relative activities of five different phytohormone signaling pathways in modulating the TSWV-thrips interaction as per Studham and MacIntosh (2012). Four of the five pathways (SA, JA, ABA and AUX) appeared to be modulated by TSWV-infection alone or in combination with thrips (**Table 2**), however plants infected with TSWV alone exhibited an overall negative effect (pathway score = −58.28) on the JA pathway. Thrips infestation on both non-infected and TSWV-infected plants activated the JA pathway, indication that the negative effect of TSWV alone on JA gene expression (**Fig. 4B**) was neutralized by the large positive effect of thrips feeding and/or oviposition on the JA pathway (pathway score = 40.02 and 56.99). Tomato plants singly-challenged by pathogen or pest exhibited an apparent negative co-regulation or cross-talk between the SA and JA pathways, however under dual challenge, both SA and JA pathways were generally up-regulated, suggesting that other factors, possibly the large ABA effect in the dual treatment (pathway score = 76.36), modulated the SA-JA crosstalk. It was also apparent that virus infection, regardless of thrips infestation, stimulated the ABA pathway and suppressed AUX pathway-associated genes, and had no apparent effect on the ET pathway even after infestation with thrips, indication that the observed thrips-only effect on the ET pathway (**Table 2**, pathway score = 40.01) was neutralized by TSWV infection (pathway score = 0.64) in the dual treatment. In total, our analysis revealed reciprocal modulation of key phytohormone pathways under dual attack. The phytohormone related genes used for this analysis are listed in **Supplementary Table 7**.

### Gene co-expression network analysis

Network analysis was performed on WGCNA-generated modules of co-expressed genes (**Supplementary Table 8**) to visualize connections among functional categories of interest (color coded) and differentially-expressed, annotated genes (**Supplementary Fig. 1**), with the knowledge that the modular structure of complex networks plays a critical role in their functionality. The result of our network analysis indicated that the gene co-expression network contains five tightly connected groups of gene sequences, whose expression is highly correlated. While photosynthesis is the predominant functional category in all of the modules, each of the groups contain genes from other functional categories, such as plant hormone-related (ABA, AUX, ET, JA, SA), protein metabolism, cell wall organization and defense functions, thus illustrating the interconnectedness among primary biological processes significantly perturbed by the virus and vector, and co-regulation of genes involved in primary metabolism (photosynthesis, cell wall organization, and protein synthesis) and defense responses.

### Reverse transcription-quantitative PCR (RT-qPCR) validation of microarray hybridization data

Six genes associated with the SA, JA, and antiviral small-RNA-mediated gene silencing pathways that were determined to be differentially-expressed by the microarray hybridization experiment were further validated by reverse transcription-quantitative PCR (RT-qPCR) (**Table 3**). We included RNA-directed RNA polymerase 1 (RDR1), a key siRNA-pathway gene involved in the amplification of virus derived siRNAs that target viral dsRNAs for degradation, to examine antiviral defense. Overall, the average relative expression ratios of SA and JA marker genes, and RDR1 in response to virus infection alone or in combination with thrips mirrored the direction (positive or negative) of expression for the microarray analyses (**Table 3**). Furthermore, pairwise comparisons of treatment averages obtained for the microarray and real-time RT-qPCR analyses revealed similar patterns among treatments (**Table 3**). The normalized abundance (NA) of TSWV nucleocapsid (N) and silencing suppressor (NSs) RNA relative to EF1 was also determined to estimate virus titer in leaf tissue. For virus infected plants (+/− thrips), there was a significant correlation (Pearson) (r = 0.996, *P* < 0.0001) between normalized abundance of N RNA and NSs RNA. Analysis of variance revealed significantly lower virus titers in leaves from TSWV-infected plants infested with *F. occidentalis* compared to plants infected with TSWV alone (**Fig. 7**) (NSs RNA: *P* = 0.006; N RNA: *P* = 0.067), indicating that thrips infestation on infected leaves for one week significantly influenced virus accumulation. There were significant correlations (n = 6) between TSWV titer and Log_2_RER for NPR1 (NSs: r = 0.941, *P* = 0.005; N: r = 0.935, *P* = 0.006), and Cathepsin D inhibitor protein (NSs: r = −0.877, *P* = 0.022; N: r = −0.897, *P* = 0.015), an indication that expression of SA- and JA-associated transcripts may be quantitatively associated with the extent of TSWV infection.

**Figure 7.**
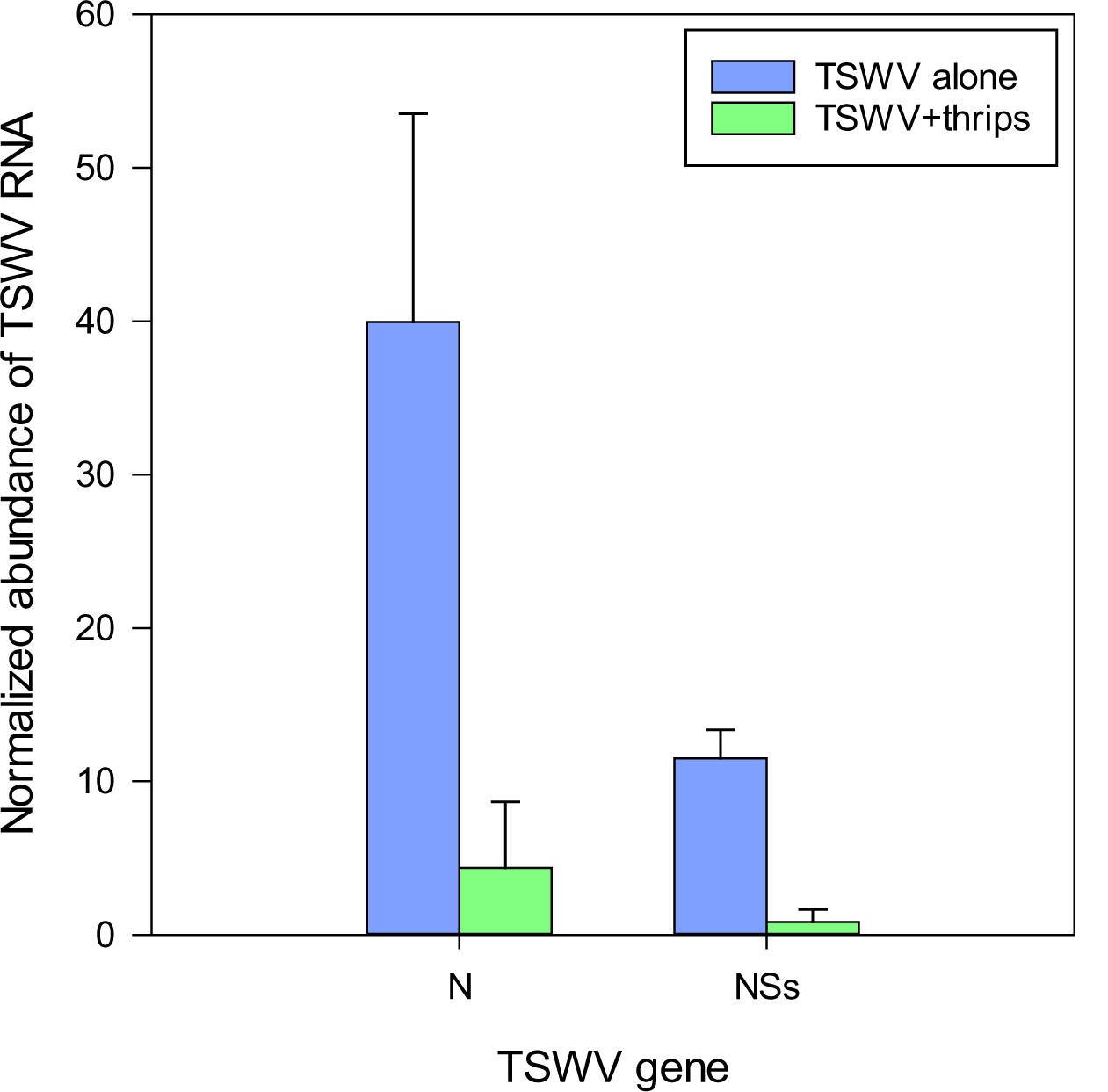
Estimation of virus titer in tomato plants systemically-infected with TSWV, three-weeks post rub-inoculation. TSWV abundance was estimated by real-time quantitative reverse transcription PCR of TSWV N and NSs RNA normalized to tomato elongation factor 1 (tEF1) RNA in the 10^th^-youngest leaf of plants infected with TSWV alone (TSWV) or infested with *Frankliniella occidentalis* females (TSWV+thrips). Each bar represents the mean (and standard error) of three biological replications (n = 3 plants) of the thrips performance/microarray experiment. Analysis of variance revealed significant differences between the TSWV and TSWV+thrips treatments [N, *P* = 0.067; NSs, *P* = 0.006].

### TSWV infection altered phytohormone levels

To determine single and combined effects of virus infection and thrips activity on the levels of the signal molecules (i.e., phytohormones), we quantified the levels of SA, JA, JA-IL, and OPDA in leaf tissues seven-days post thrips release. Same-age leaflets that were immediately basal (older) to the leaves harvested for microarray analysis were used for phytohormone analysis. TSWV infection alone or in combination with thrips enhanced SA content in tomato leaves (**Table 4**). Thrips feeding had no apparent effect on any of the phytohormones at the time of sampling; however, the combined effect of TSWV and thrips resulted in significantly higher leaf contents of JA. There were no apparent differences among treatments with regards to JA-IL or OPDA at the time of sampling (**Table 4**).

**Table 4.** Phytohormone content expressed as Log_10_(ng analyte g fresh weight^-1^) in leaflets samples from tomato plants systemically-infected with TSWV and/or infested with thrips one week-post thrips release. Values represent mean ± standard deviation (n = 4 plants). Different letters indicate statistical difference between treatments at *P* ≤ 0.05.

| **Phytohormone** | **Mock** | **TSWV** | **Thrips** | **TSWV + thrips** |
| --- | --- | --- | --- | --- |
| Salicylic acid | 1.72 ± 0.12b | 3.15 ± 0.18a | 1.85 ± 0.13b | 2.90 ± 0.22a |
| Jasmonic acid | 0.16 ± 0.06b | 0.29 ± 0.14b | 0.25 ± 0.11b | 0.67 ± 0.14a |
| JA-Isoleucine | 0.05 ± 0.48a | 0.26 ± 0.36a | 0.24 ± 0.50a | 0.92 ± 0.36a |
| OPDA (12-oxo-phytodienoic acid) | 0.60 ± 0.39a | 1.15 ± 0.34a | 0.89 ± 0.34a | 1.33 ± 0.31a |

### TSWV infection increased total free amino acid content

As a measure of plant quality to the insect vector, we measured total free amino acid content to determine if TSWV infection altered the nutritional status of the host. There was a significant effect of time (biological replicate) (*F*=10.87, df=2, *P* = 0.0002) and treatment (*F*=4.25, df=3, *P* = 0.01) but not the interaction term (*F*=1.11, df=6, *P* = 0.37) on the total free amino acid content. Overall, TSWV infection alone and in combination with thrips feeding harbored greater total free amino acid content (Mean ± SE: 288.64 ± 110.87 and 290.12 ± 64.85, respectively) compared to mock-inoculated and thrips –fed plants (Mean ± SE: 164.03 ± 20.56 and 181.55 ± 25.23, respectively) (**Fig. 8**).

**Fig. 8.**
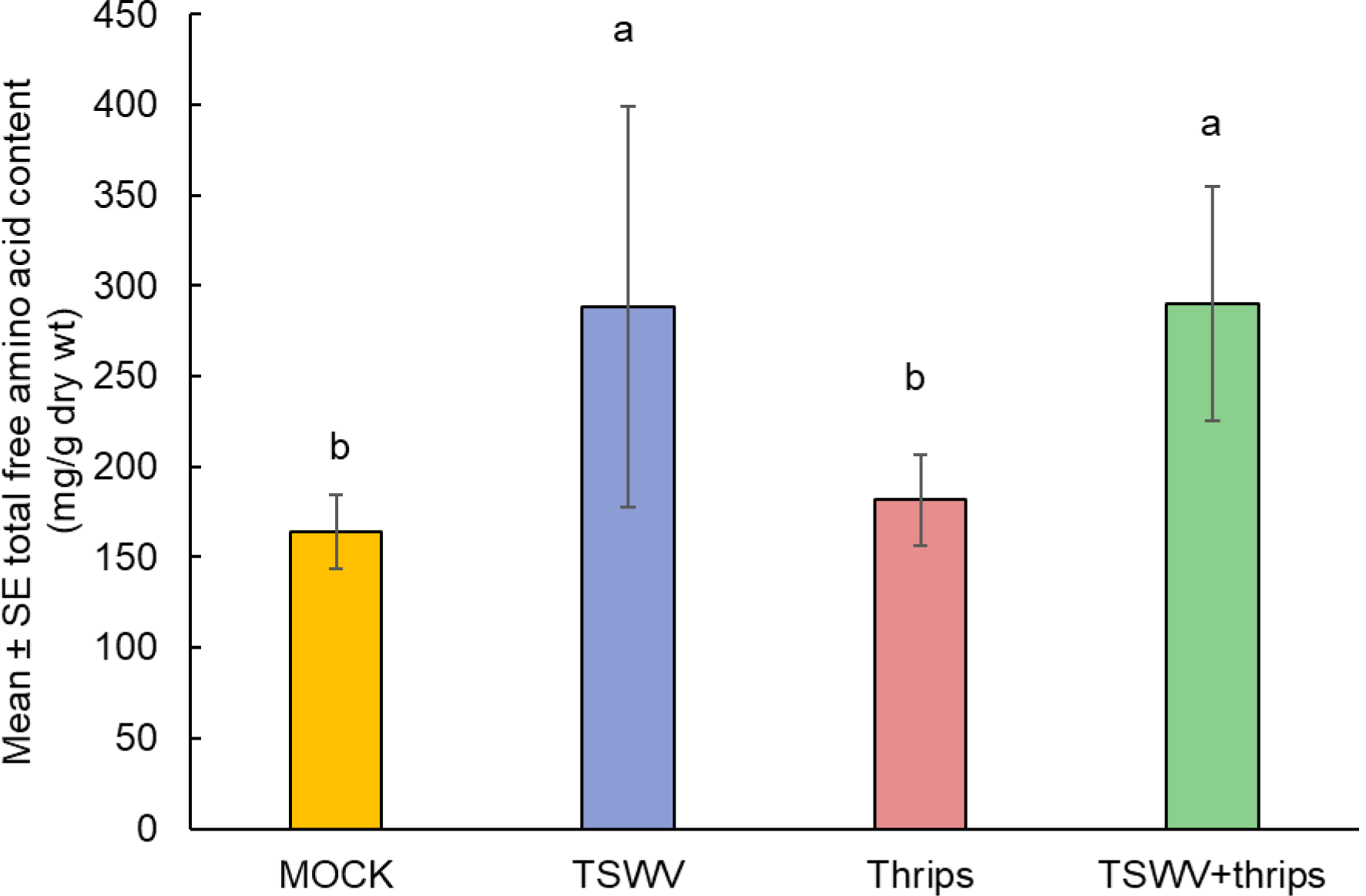
TSWV infection increased total free amino acid content in tomato plants. Tomato plants were systemically-infected with TSWV and/or infested with *F. occidentalis*. Each bar represents the mean± standard error of n4 plants per biological replicate. Different letters indicate significant differences between treatments at *P<*0.05.

## DISCUSSION

Plants encounter and respond to multiple and often co-occurring biotic stresses such as pathogens and insect herbivores in a myriad of ways. Moreover, plant pathogens and insects that share the same host are likely to interact and these interactions may have positive, negative or neutral consequences (Belliure et al., 2005; Belliure et al., 2010; Thaler et al., 2010; Pan et al., 2013; Eigenbrode et al., 2018). For instance, previous research by us and others showed that a plant virus, TSWV enhanced survival and oviposition of a non-vector herbivore, spider mites (Belliure et al., 2010; Nachappa et al., 2013). In this study, we explored plant responses that underlie the impact of TSWV on its thrips vector, *F. occidentalis*. While previous research identified plant defense response as a key molecular mechanism through which TSWV affects its vector (Abe et al., 2012; Wu et al., 2019), other pathways could be involved (Nachappa et al., 2013). Here, we demonstrate that virus infection alters the expression of a large set of coordinated networks of regulatory genes controlling primary metabolic pathways and defense responses thereby rendering virus-infected plants more suitable hosts for its insect vectors. To our knowledge, this is the first study to identify global transcriptional networks that modulate TSWV-thrips interaction in tomato and provides information on key pathway players.

Orthotospoviruses depend solely on the thrips vector for transmission (Rotenberg and Whitfield, 2018). Several studies documenting changes in thrips settling behavior/host preference, feeding behavior and/or performance due to virus infection show the possibility of positive, negative or neutral outcomes (reviewed in Eigenbrode et al. (2018). Positive benefits to a vector are predominantly associated with indirect manipulation of the vector via the host, whereas negative effects are associated with direct manipulation of the vector after acquisition of the virus (Belliure et al., 2005; Shrestha et al., 2012). In the current study, we found that tomato plants infected with TSWV enhanced performance of thrips compared to healthy or mock-inoculated plants. Previous studies have shown that virus infection increases free amino acid content in infected plants (Markkula and Laurema, 1964; Shrestha et al., 2012), which is known to impact vitellogenesis in thrips and also serve as nutrients for the developing eggs (Klowden, 2013). In agreement with these results, we found that virus infection alone or in combination with thrips resulted in 2.5 and 1.9 times greater total free amino acid content compared to mock-inoculated plants. This suggests that the increased population of thrips is influenced by the increased concentrations of total free amino acids in TSWV-infected plants. In choice assays, we found that TSWV infection increased attractiveness of the host plant for thrips vectors. Recently, Wu et al. (2019) showed that TSWV infection enhances plant attractiveness to the thrips vector by suppressing synthesis of volatile plant terpenes, which is known to repel herbivores including thrips. There were two oligos related to terpene synthesis in the tomato GeneChip^®^, monoterpene synthase 1 and sesquiterpene synthase 1, but there were not differentially-expressed (*P*<0.05, fold change >2) between the treatments (data not shown). Taken together, increased aggregation and population growth of thrips on virus-infected plants can potentially increase the number of viruliferous vectors, which is expected to increase virus spread. Importantly, the attraction of adult female thrips to TSWV-infected plants would promote oviposition on virus infected plants. Larval thrips, the only life stage that can effectively acquire TSWV, that emerge would be able to transmit later in development thus completing a TSWV transmission cycle.

Plants infected with viruses are often found to be more suitable hosts for insect vectors than uninfected plants (Belliure et al., 2005; Blanc and Michalakis, 2016; Mauck, 2016; Eigenbrode et al., 2018). Until now, most studies have focused on virus-induced suppression of anti-herbivore defenses (Thaler et al., 2010; Abe et al., 2012; Casteel et al., 2012; Mauck et al., 2019; Wu et al., 2019). A recent study found that plants mount several layers of defense to resist attack from TSWV infection (reviewed in Zhu et al. (2019). Consistent with these findings, our study demonstrates that virus infection up-regulated a suite of genes related to plant innate immunity and defense response. In the TSWV-thrips interaction, virus infection has been shown to increase the anti-pathogen response (SA-related defenses) which suppresses the anti-herbivore response (JA-related defenses) by exploiting the antagonistic crosstalk between SA-JA plant defenses. This attracts and benefits the vector thrips compared to uninfected plants (Abe et al., 2012). Similarly, we found that virus infection induced a majority of SA-regulated genes and SA content in the plants, but only 29% of the JA-related responses in virus-infected plants were repressed in these plants, which suggests that the inhibition by SA of JA responsive genes is transient. SA-mediated suppression of JA-responsive gene expression is thought to mainly occur downstream of the JA biosynthesis pathway (Leon-Reyes et al., 2010). This may explain the down-regulation of JA-inducible genes such as wound-induced proteinase inhibitors, and the lack of down-regulation in JA biosynthesis genes such as *OPR3*, *LOX* AND *AOS* in TSWV-infected plants. Mechanical wounding and insect feeding activate JA-regulated wound response genes such as proteinase inhibitors (Green and Ryan, 1972; Walling, 2000; Wasternack et al., 2006). Moreover, it is likely that timing and magnitude of responses play a major role in orchestrating SA-JA antagonism. For example, Abe et al. (2012) found that the magnitude of suppression of JA-related genes was reduced at 7-d compared to 14-d post-TSWV infection. In the current study, gene expression was measured 3 weeks post-virus-infection, which may be one reason for attenuation of plant responses. Moreover, the host plants were different in the two studies, tomato (current study) versus *Arabidopsis* (Abe et al., 2012), which may in part explain the inconsistencies between the two studies. Further experimentation utilizing tomato mutants of the SA- and JA-signaling pathways would be useful to explore this result. Koornneef et al. (2008) showed that the antagonistic effect of SA on JA signaling was evident when SA was applied simultaneously with MeJA; however, when SA was applied more than 30 h prior to the onset of the JA response, SA-mediated suppression of JA was not observed. These results suggest that SA-JA crosstalk is transient and depends on the timing and magnitude of elicitation. Given the dynamic nature of TSWV infection and symptom development (chlorosis, stunting and wilt), it is crucial to analyze gene expression profiles during early versus late stages of infection or disease development as it relates to vector performance. While the roles of SA and JA/ET in plant defenses are well-established, these hormones also affect a myriad of development processes such as growth repression, flower development and fertility (JA), flowering time (SA), and germination, senescence and fruit ripening (ET) that may influence outcomes of virus infection. This suggests an intimate interplay exists between phytohormone regulation and primary metabolism (Reviewed in (Rojas et al., 2014; Berens et al., 2017).

Other phytohormones such as ET, ABA, auxins, brassinosteroids, cytokinins and gibberellins also play in role in plant-pathogen interactions (De Bruyne et al., 2014; Shigenaga and Argueso, 2016; Yue et al., 2016; Berens et al., 2017; Yang et al., 2019). In the current study, we found that virus infection alone and in combination with thrips up-regulated ABA-related genes and down-regulated auxin-related genes. In general, ABA acts antagonistically with SA, hence it may be beneficial for the virus to induces ABA genes, resulting in suppression of anti-pathogen or SA-mediated defenses (Mauch-Mani and Mauch, 2005; Asselbergh et al., 2008). The role of ABA in response to insect feeding is not clear; however studies have shown that aphids induce ABA as a decoy strategy to suppress SA- and JA-related defenses (Studham and MacIntosh, 2013; Nachappa et al., 2016). Thrips feeding down-regulated ABA-related genes. Auxin regulates plant growth and development, and is also involved in stress responses (Reviewed in (Kazan and Manners, 2009). Auxin can affect disease outcomes directly and indirectly. Direct interaction may involve interaction with other phytohormones including SA, JA, and ET. For example, auxin suppresses immunity against the bacterial pathogen *Pseudomonas syringae* via SA suppression (Wang et al., 2007). Indirect effects of auxin may involve changes in plant growth and development and thereby outcomes of disease resistance. Virus-infected plants often show developmental abnormalities such as stunting and leaf curling, which resemble auxin mutants leading to the conclusion that virus infection alters host auxin signaling. We found that TSWV infection alone and the dual treatment suppressed auxin genes potentially resulting in TSWV-related developmental abnormalities; however, the impact of such developmental alterations on the insect vector is not known.

Although at different time points post-infection, similar trends documented in the transcriptome profiles of tomato and *Arabidopsis* during TSWV were consistent with our findings (Catoni et al., 2009; Padmanabhan et al., 2019; Xu et al., 2020). We collected plant samples after thrips infestation (3 weeks after virus inoculation) but the other published work suggests that at the time of first thrips deposition on plants, 2 weeks after TSWV-inoculation, these important defense responses and plant barriers were compromised and promoted thrips performance on plants infected with TSWV. Notably, we found that virus infection alone and the combined treatment of virus and thrips down-regulated majority of the photosynthesis-related genes such as ribulose bisphosphate carboxylase, ATP binding protein, chaperone, and chlorophyll a/b-binding proteins. Although considerable progress has been made in understanding the plant defense responses, very little is known about the role of primary metabolic pathways associated with plant growth and development in regulating defense responses. It is hypothesized that the energy saved by down regulation of primary metabolism specifically photosynthesis and chlorophyll biosynthesis is diverted and used for defense responses (Less et al., 2011; Rojas et al., 2014). Moreover, the chloroplast, houses several important steps in the synthesis of phytohormones involved in defense, such as SA, JA, and ABA (Reviewed in (Kangasjärvi et al., 2012)). Hence, chloroplast-related proteins potentially crosstalk with defense-related proteins. The chloroplast is also the engine of plant growth and plays a crucial role in symptom development. For example, the development of chlorotic or yellowing symptoms of virus-infected plants is likely due to suppression of chloroplast pigment genes. Previous research found that TSWV-infected lettuce plants were more attractive for thrips vectors compared to healthy plants because of the yellow color of the infected plants (Yudin et al., 1987). Catoni et al. (2009) also found that genes related to photosynthesis were suppressed in tomato leaves 14 dpi. More recently, Xu et al. (2020) studied gene expression in response to TSWV infection in *Arabidopsis* across different development stages (9, 12 and 15 dpi). They too found that genes such as rubisco and Chaperonin and others involved in the photosynthesis pathway were largely suppressed in response to TSWV infection.

In this study, we found that virus infection altered genes related to cell wall organization or biogenesis. Most cell-wall-modification enzymes such as xyloglucan endochitinase, endotransglycosylase/hydrolases and pectinesterase expression levels were decreased in virus-infected plants. In contrast, cell wall degradation genes such as expansin, some cellulases and beta-1, 3-glucanases were upregulated. In agreement with our findings, Xu et al. (2020) found that cellulose synthesis genes were down-regulated whereas cellulases and beta-1,3-glucanases were up-regulated to promote cell wall degradation. Interestingly, beta-1, 3-glucanase is a SA-inducible PR gene that results in SA accumulation during pathogen attack. In a study on transcriptome analysis of a TSWV resistant tomato line (*Sw-7*) and susceptible line challenged with TSWV infection, the authors found cell wall-related genes to be down-regulated in the resistant line, *Sw-7* and only three out of the five pectinesterases were up-regulated (Padmanabhan et al., 2019). The authors also reported up-regulation of pathogenesis-related proteins, *PR-1* and *PR-5* (osmotin) that modulate callose and lignin deposition in the cell wall leading to restricted virus movement in the resistant line. Indeed, our study also found increased expression of *PR-1* and *PR-5* in virus-infected plants. These findings provide evidence supporting the dual regulatory role of cell wall organization genes in plant structure and defense. It is possible that suppression of cell wall genes renders the plant more susceptible to the penetration of mouthparts of thrips vectors.

Another functional category of interest was protein synthesis and degradation. Plants are known to accelerate protein metabolism to resist virus infection and spread (Verchot, 2016). In the current study, genes related to protein synthesis and degradation were induced in response to TSWV infection alone and in combination with thrips feeding. Specifically, genes associated with ribosome 60S or 40S subunits were induced as were genes associated with protein degradation such as those that encode 26S proteasome and in the ubiquitination pathway. These results are in agreement with a previous report of increased up-regulation of protein degradation genes during TSWV infection (Xu et al., 2020). Analysis of total free amino acids revealed increased AA content in TSWV-infected plants compared to mock and thrips infested plant. This increase may in part be due to increased protein metabolism in virus-infected plants. It is likely that induction or suppression of specific amino acids is involved in plant defense against pathogens and insects. However, since we did not analyze individual amino acids, we cannot make conclusions about the role of amino acids in defense. There are reports of the involvement of particular amino acids in defense. For example, the *lht1* (lysine histidine transporter 1) mutant of *Arabidopsis* has significantly reduced contents of glutamine, alanine, and proline in comparison with wild-type plants and showed enhanced resistance to diverse bacterial, fungal, and oomycete pathogens (Liu et al., 2010)

The positive effect of TSWV on *F. occidentalis* described here is consistent with the work of other researchers, but we cannot overlook the broad acting effects of TSWV on non-vector and other trophic levels. Indeed, previous research showed that TSWV infection had positive effects on non-vectors as well (Belliure et al., 2010; Nachappa et al., 2013) by modulating the same pathways namely defense response, photosynthesis, cell wall metabolism and protein metabolism (Nachappa et al., 2013). Hence, changes in these key pathways may also be exploited by non-vector herbivores. Interestingly, one study found that TSWV-infected female thrips were more predaceous on spider mite eggs compared to uninfected thrips (Stafford-Banks et al., 2014). While this behavior is unlikely to increase virus transmission directly, TSWV infection appears to indirectly enhance fitness of thrips vector by improving oviposition of the spider mite prey. In contrast, Pan et al. (2013) found that TSWV infection decreased fitness of whiteflies on peppers. Further research is required to determine whether these mechanisms are different from those identified in our study. Virus infection can also alter plant volatiles that can have consequences for both vector and non-vector herbivores. For example, significantly more aphids settled onto barley yellow dwarf virus-infected than non-infected plants (Jiménez-Martínez et al., 2004). In contrast, fungus gnat adults preferred non-infected plants compared to white clover mosaic virus-infected plants based on their volatile blends (van Molken et al., 2012). These results suggest that the consequences of virus infection on vector and non-vectors and the third trophic level depends on the specific ecological setting, i.e. the species involved, the order in which they attack, the time span of the interaction, the frequency and spatial scales at which pathogens and herbivores co-occur (Roossinck, 2011; van Molken et al., 2012).

In the current study, TSWV appeared to be the prominent driver of plant responses, with some modulation by the thrips vector. With a few specific exceptions, the combination of TSWV and thrips mirrored global gene expression patterns of TSWV infection alone, which suggests that TSWV infection has a significant effect on plant physiology compared to thrips. However, this was likely due, in part, to the length of time allowed for virus accumulation and symptom development (two weeks) prior to release of female thrips on these plants. Interestingly, we found that thrips – on infected plants for only one week prior to sampling – had a negative effect on virus titer as measured by real time qRT-PCR using two viral genes, TSWV-N and NSs. The expression of canonical virus-activated pathways was similar for TSWV alone and the dual treatment leading us to hypothesize that i) the dual attack may further compromise plant health and thus creates a less robust host, or ii) the presence of the insect vector alters the virus composition in the plant host through a yet unidentified pathway or mechanism. Further exploration of this phenomenon is warranted and it will be interesting to determine if thrips perception by the plant alters the virus in a way to promote acquisition similar to transmission morphs described for non-circulative viruses (Gutiérrez et al., 2013; Martinière et al., 2013; Berthelot et al., 2019).

## Acknowledgements

This work was supported by a seed grant from The Ecological Genomics Institute at Kansas State University. We thank Alexandria Leach-Kieffaber and Vamsi J. Nalam for their assistance with the RT-qPCR experiments.

## Supplemental Material

**Supplementary Fig. 1.** Weighted gene co-expression network (WGCNA) plotted using the ‘layout_with_fr’ layout in igraph. Each node represents a probe set id (sequence) from the gene chip. The five clusters of nodes following leading eigen community detection are illustrated by the different colored nodes. The membership of nodes in each community is provided in Supplementary Table 8.

**Supplementary Table 1.** List of differentially-expressed genes (fold change ≥2.0 and *P*< 0.05) in tomato plants systemically-infected with TSWV.

**Supplementary Table 2.** List of differentially-expressed genes (fold change ≥2.0 and *P*< 0.05) in tomato plants infested with *F. occidentalis*.

**Supplementary Table 3.** List of differentially-expressed genes (fold change ≥2.0 and *P*< 0.05) in tomato plants systemically-infected with TSWV and infested with *F. occidentalis*.

**Supplementary Table 4.** List of differentially-expressed genes associated with plant defenses and phytohormone pathways.

**Supplementary Table 5.** List of differentially-expressed genes associated with photosynthesis, cell wall organization and protein metabolism.

**Supplementary Table 6.** List of differentially-expressed genes associated with specific clusters in the heatmap or hierarchical cluster analysis.

**Supplementary Table 7.** List of differentially-expressed genes used to generate phytohormone pathway scores.

**Supplementary Table 8.** List of differentially-expressed genes associated with each cluster of nodes or community in the WGCNA analysis and gene co-expression network.

## Notes

### Competing Interest Statement

The authors have declared no competing interest.

