## Supplementary figures and images for "Tomato spotted wilt virus benefits its thrips vector by modulating metabolic and plant defense pathways in tomato"

### Supplemental Fig 1

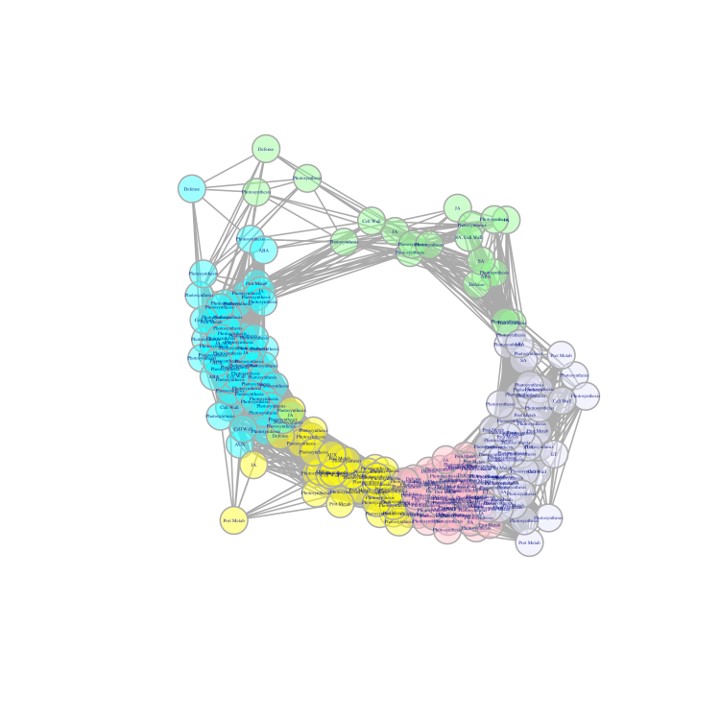
